# O-GlcNAc transferase regulates H_2_O_2_ production via p38 MAPK

**DOI:** 10.64898/2026.05.27.728188

**Authors:** Luke I Jones, Shia Vang, Hannah J McIntire-Ray, Holly A Petersen, Angela N Morales, Viviana E Acevedo Rua, Joshua C Anderson, Andres J Gonzalez Coba, Stefanie Krick, Jarrod W Barnes

## Abstract

i.

Idiopathic pulmonary fibrosis (IPF) is a progressive interstitial lung disease characterized by augmented transforming growth factor-β (TGF-β) signaling leading to excessive extracellular matrix (ECM) deposition. The fibroblast-to-myofibroblast-transition (FMT) and metabolic reprogramming of lung fibroblasts (HLFs) are essential to IPF pathogenesis, yet the connection between nutrient metabolism and fibrogenesis remains poorly defined. The O-linked N-acetylglucosamine (O-GlcNAc) transferase (OGT) is a nutrient-sensitive enzyme that adds O-GlcNAc moieties to substrates. We previously showed that loss of OGT reverses bleomycin-induced pulmonary fibrosis in mice. Here, using unbiased kinomics, we show that pharmacologic inhibition of OGT suppressed non-canonical TGF-β-induced mitogen-activated protein kinase (MAPK) signaling. Molecular confirmation revealed that TGF-β-induced phosphorylation of p38, but not ERK or JNK, was reduced by OGT blockade. Furthermore, p38 itself was O-GlcNAc-modified, which enhanced its phosphorylation and promoted downstream phosphorylation of the NADPH oxidase subunit, p47^phox^. Inhibition of OGT, p38, or p47^phox^ reduced reactive oxygen species (ROS) in HLFs, revealing a previously unknown role of OGT-p38-p47^phox^ signaling in ROS production. Collectively, this work establishes that O-GlcNAc-modified p38 enhances p47^phox^-dependent H_2_O_2_ production.

**Highlights:**

1. Using PamChip STK arrays, we show that OGT inhibition causes broad kinomic remodeling, including suppression of non-canonical TGF-β MAPKs and multiple CDKs.
2. OGT blockade selectively attenuates p38 phosphorylation, despite TGF-β-induced substrate redundancy with ERK and JNK.
3. We provide evidence that p38 MAPK undergoes O-GlcNAcylation in human lung fibroblasts, a modification not previously reported.
4. The study identifies a new signaling axis where O-GlcNAc modification of p38 modulates the phosphorylation of p47^phox^, therefore regulating NOX-dependent H_2_O_2_ production.
5. Blocking OGT or inhibiting p38/p47^phox^ dramatically reduces TGF-β-driven H_2_O_2_ production in human lung fibroblasts.

**Graphical Abstract:** 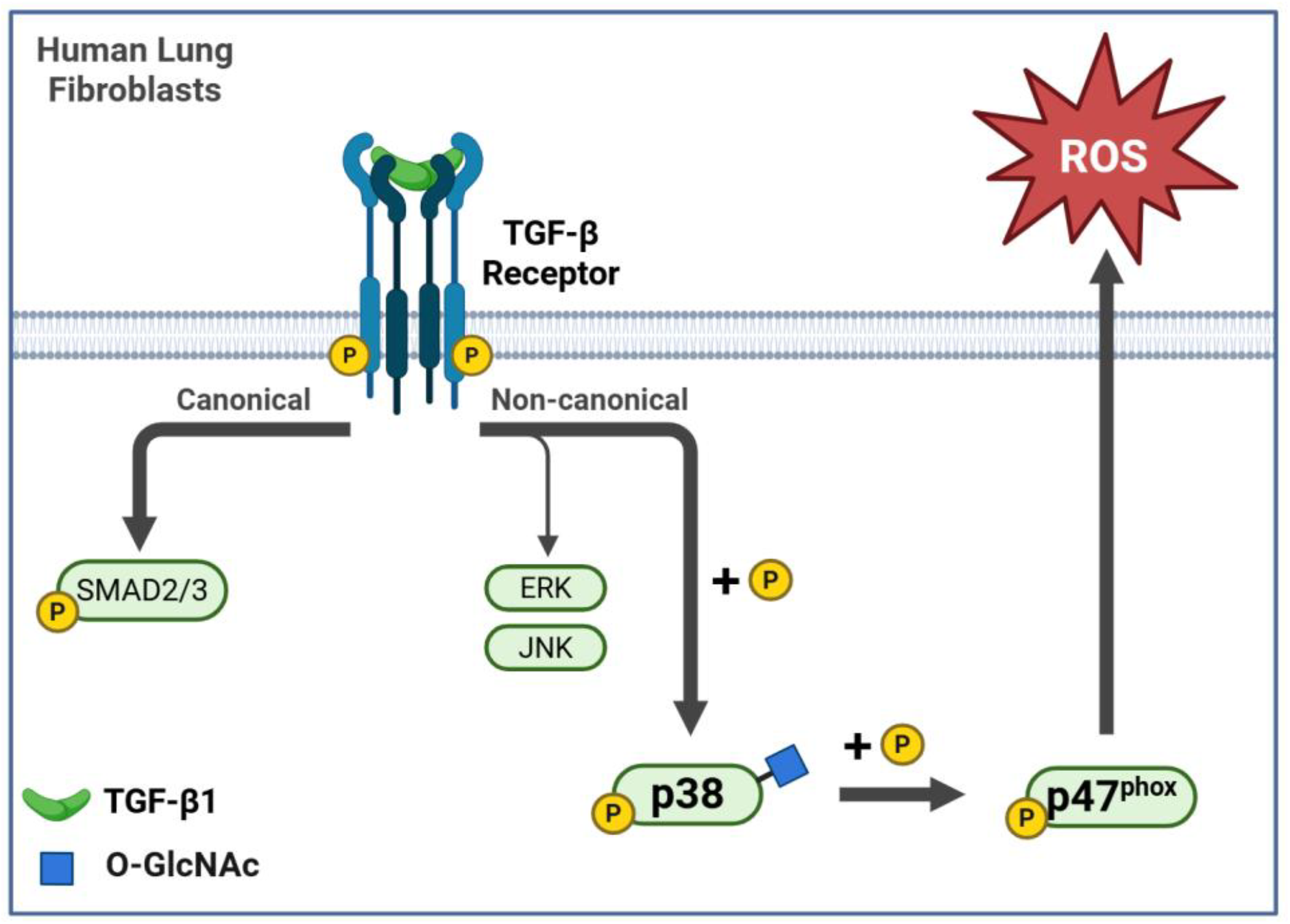

## ii. INTRODUCTION

Idiopathic pulmonary fibrosis (IPF) is the most common interstitial lung disease affecting approximately 3 million people worldwide^1, 2^. IPF is a chronic, progressive disorder characterized by irreversible scarring of the lung. Excessive fibrosis leads to destruction of alveolar architecture, resulting in impaired gas exchange and respiratory failure^3, 4^. Fibrosis is a physiological response to lung injury and is defined by the excessive deposition of extracellular matrix (ECM) proteins, including collagen, fibronectin, and elastin^5-8^. Although the etiology of IPF remains unknown, fibroblasts and myofibroblasts are recognized contributing effector cells driving fibrogenesis^6, 9^.

Without lung transplant, there is currently no known cure for IPF. Standard treatment includes three FDA-approved therapies, pirfenidone, nintedanib, and nerandomilast, which inhibit profibrotic signaling pathways to reduce fibroblast proliferation, decrease ECM deposition, and slow the rate of decline in forced vital capacity (FVC)^10-14^. While these therapies modestly improve disease progression, IPF remains associated with a poor prognosis, with median survival ranging from 2-5 years post-diagnosis. IPF pathogenesis involves multiple aberrant cellular processes, including chronic inflammation, dysregulated growth factor signaling, impaired ECM clearance, metabolic dysregulation, and altered cell survival. Therefore, a more comprehensive understanding of the molecular mechanisms regulating these pathologic processes is needed to identify more effective therapeutic strategies.

The transforming growth factor–β (TGF-β) superfamily is a well-established profibrotic signaling pathway in pulmonary fibrosis. Notably, canonical SMAD3 signaling is well documented for its role in IPF fibrogenesis^15-18^. In addition to SMAD-dependent pathways, SMAD-independent (non-canonical) signaling pathways, including members of the mitogen-activated protein kinase (MAPK) family (e.g. ERK, p38, JNK) have also been implicated in IPF pathogenesis through inflammation, myofibroblast differentiation and proliferation, production of reactive oxygen species (ROS), and resistance to apoptosis^19-25^.

Metabolic reprogramming of activated fibroblasts and myofibroblasts is required to support the high energetic and biosynthetic demands associated with sustained proliferation and ECM production^26, 27^. The O-linked N-acetylglucosamine (O-GlcNAc) transferase (OGT) functions as a metabolic stress and nutrient sensor that catalyzes the transfer of GlcNAc moieties to serine/threonine residues of proteins. Both OGT and O-GlcNAcase (OGA), the hydrolase that removes O-GlcNAc, have been reported to control signal transduction, transcriptional activity, and cellular adaptation through the dynamic regulation of O-GlcNAc^28^. Emerging evidence demonstrates extensive crosstalk between O-GlcNAc and phosphorylation, whereby these modifications can occur reciprocally or cooperatively on proteins to alter kinase activation, substrate affinity, protein stability, and downstream signaling pathways^29-32^. In particular, MAPK signaling is highly dependent on tightly regulated phosphorylation kinetics; however, whether O-GlcNAcylation contributes to the regulation of non-canonical TGF-β MAPK signaling in human lung fibroblasts remains unknown.

We previously identified that pharmacologic and genetic inhibition of OGT attenuates canonical TGF-β signaling through O-GlcNAc modification of SMAD3, reversing hallmarks of fibrosis in human lung fibroblasts and a murine model of pulmonary fibrosis^33-35^. OGT may however regulate other signaling pathways relevant to fibrogenesis. Therefore, we hypothesized that O-GlcNAc transfer may broadly coordinate phosphorylation-dependent kinase networks involved in fibroblast activation.

Utilizing an unbiased kinomics platform, this study assessed pharmacologic inhibition of OGT on TGF-β-induced kinase activity in human lung fibroblasts. Inhibition of OGT caused robust differences in kinase activity in HLFs, including the suppression of p38 MAPK kinase activity, which is important in fibrogenesis^36^. We demonstrate that p38 MAPK is O-GlcNAc-modified, which attenuates its phosphorylation and subsequently reduces H_2_O_2_ production via p47^phox^. Altogether, these experiments demonstrate a role for the OGT/O-GlcNAc axis in coordinating phosphorylation-dependent signaling networks in HLFs, specifically the non-canonical TGF-β MAPK axis.

## iii. RESULTS

### OGT inhibition promotes kinomic changes human lung fibroblasts

We previously identified that pharmacologic and genetic inhibition of OGT attenuates canonical TGF-β signaling and reverses fibrosis in HLFs and a murine model of IPF^33^. Using PamChip®, a high-sensitivity, cell-free serine-threonine kinase (STK) array, we evaluated the impact of OGT inhibition and TGF-β stimulation in HLFs on the phosphorylation of known STK peptides (**Fig 1a**). Differential peptide phosphorylation was observed after acute TGF-β stimulation with and without pre-treatment with an OGT inhibitor, OSMI-1 (**Fig 2a**). Using BioNavigator v6.2 with the UpKin PamApp, upstream kinases were predicted based on kinase consensus sequences and substrate phosphorylation on the PamChip® (**Fig 2a**). A volcano distribution of TGF-β stimulated mean kinase statistics predictions confirmed enrichment of the non-canonical TGF-β MAP kinases, p38, ERK, and JNK (**Fig 2b**). Interestingly, OSMI-1 pretreatment prevented acute TGF-β stimulation of these kinases in HLFs (**Fig 2c**). These findings indicate that blocking O-GlcNAc transfer via OGT inhibition attenuates non-canonical TGF-β kinase activity.

**Figure 1.**
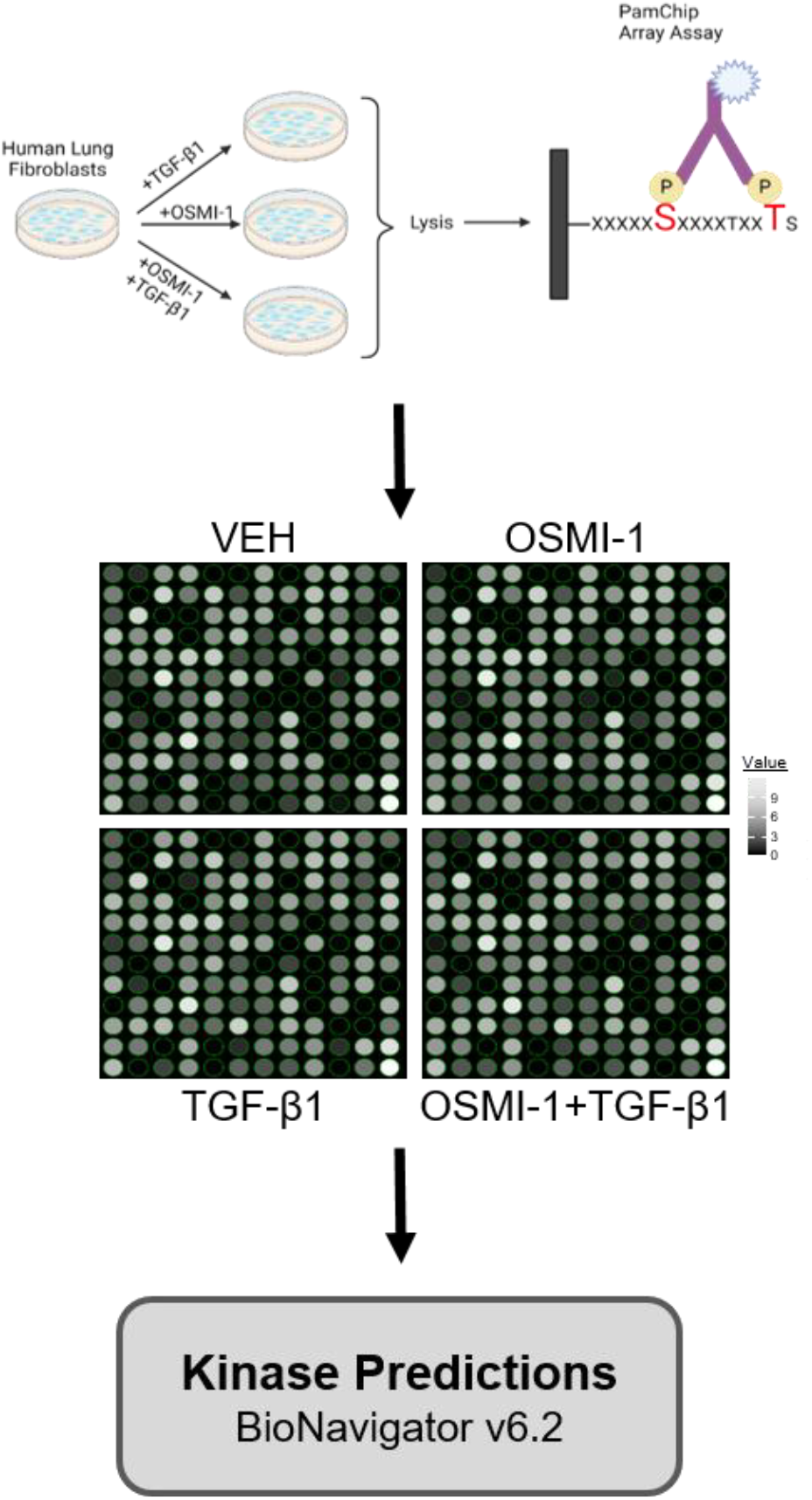
Overview of unbiased kinomics approach in human lung fibroblasts (HLFs). HLFs were serum starved overnight followed by 2-hour pretreatment with vehicle (DMSO) or OSMI-1 (25 μM). Immediately following, all groups were stimulated with or without rhTGF-β1 (5 ng/mL) for 30 minutes. HLF whole cell lysates were loaded onto a PamChip® Array for kinomics analysis to measure phosphorylation of peptide substrates. Upstream kinases of each PamChip® substrate were predicted using BioNavigator v6.2.

**Figure 2.**
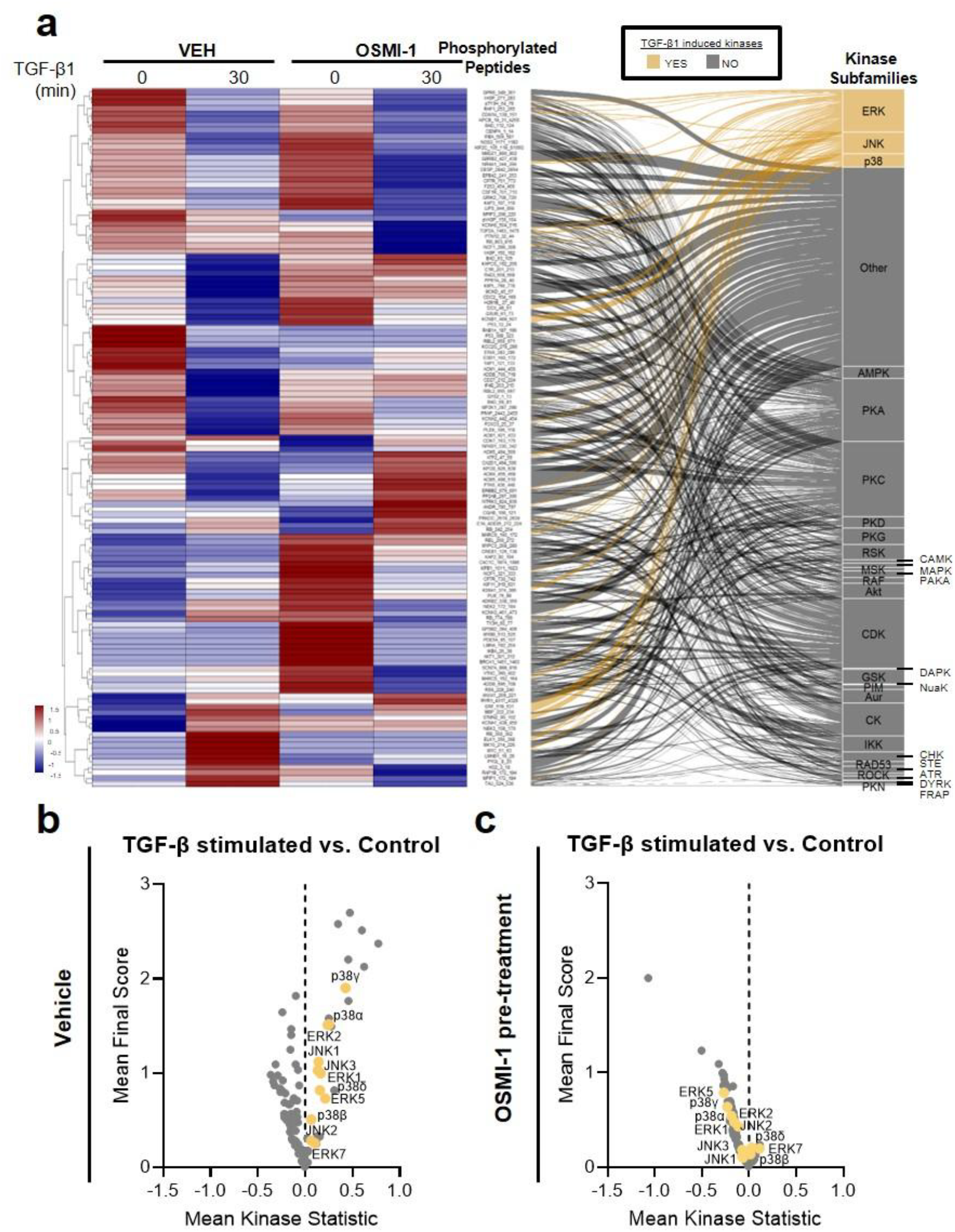
OGT inhibition prevents TGF-β-induced non-canonical MAPK activity. **A)** Heat map of differentially phosphorylated peptides from HLF pretreatment with or without OSMI-1 and/or TGF-β1 as in **Fig 1**. Upstream kinases of each phosphorylated peptide were predicted with BioNavigator v6.2. Associations between phosphorylated peptides and predicted kinase subfamilies are shown via with an alluvial plot; TGF-β stimulated kinases are indicated in gold. **B)** Volcano distribution of TGF-β stimulated kinases, reported as mean kinase statistic by final score (see *Methods*) **C)** Volcano distribution of TGF-β stimulated kinases following OSMI-1 pretreatment (25μM; 2hrs). Non-canonical TGF-β MAP kinases are indicated in gold.

### OGT is critical for p38 MAPK phosphorylation in HLFs

There is significant substrate redundancy among p38, ERK, and JNK kinases. It remains to be determined whether OGT blockade globally affects this family or whether differential, kinase-specific effects exists. As the activity of ERK, JNK, and p38 MAPK is dependent on their phosphorylation status, we sought to determine whether loss of O-GlcNAc transfer via OSMI-1 affected TGF-β-induced phosphorylation of these kinases. Following OGT inhibition in HLFs, a global reduction in phosphorylated serine/threonine residues was observed (**Fig 3b**). This effect was interestingly indiscriminant of TGF-β1 stimulation. Under the same conditions, both JNK and ERK phosphorylation were unaffected at baseline and following acute TGF-β stimulation (**Fig 3c,d**). Conversely, TGF-β1 stimulation promoted robust p38 MAPK phosphorylation at the same time point, which was attenuated by OSMI-1 pretreatment (**Fig 3e**). This finding was not only limited to acute TGF-β stimulation but was maintained following prolonged (15 hours) ligand exposure (**Fig 3f**). These data identify OGT as a selective regulator of p38 MAPK phosphorylation, without affecting JNK or ERK signaling, and provide a mechanistic framework whereby loss of OGT attenuates p38 MAPK activation and diminishes its downstream kinase activity.

**Figure 3.**
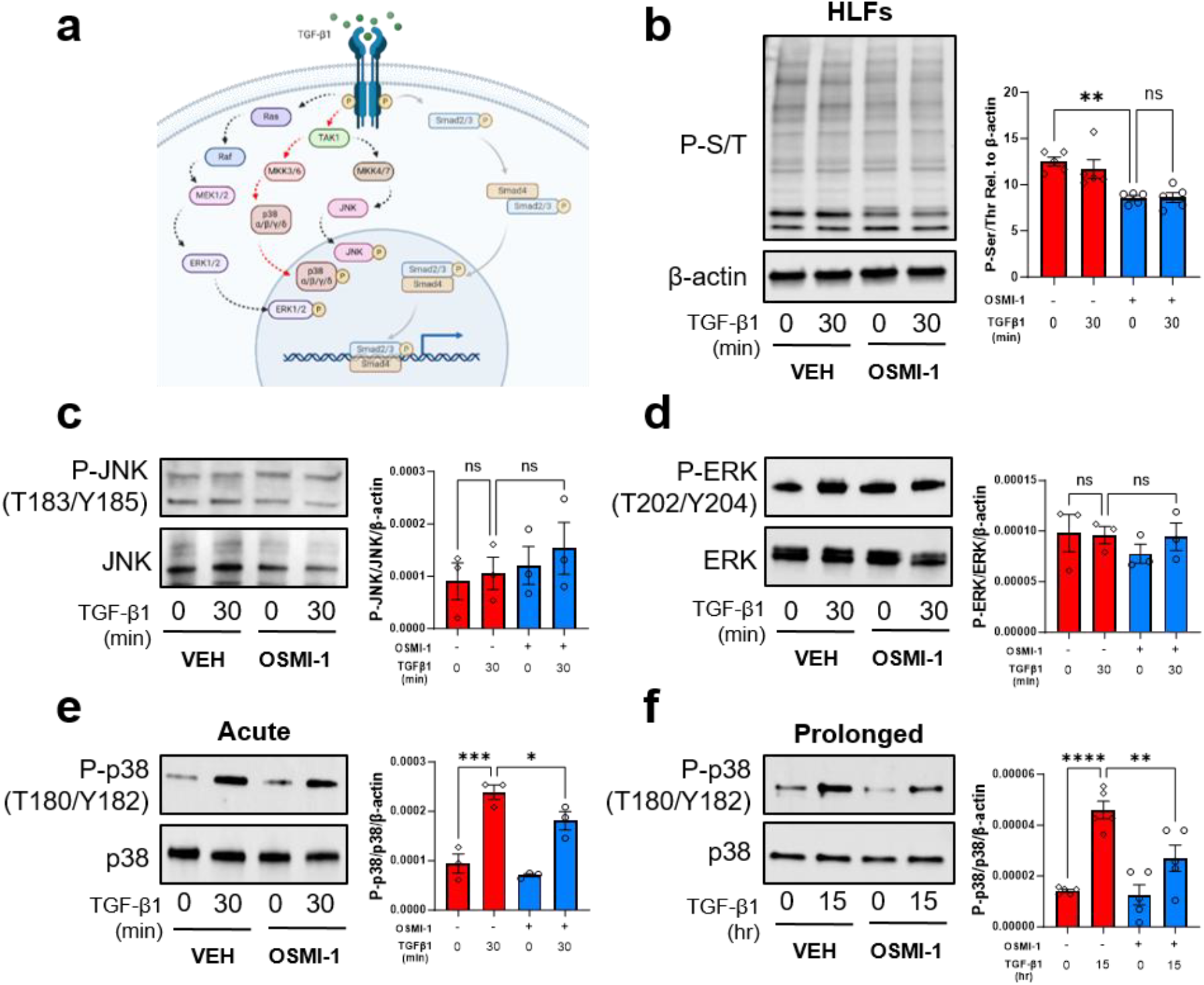
TGF-β-induced phosphorylation of p38 MAPK is regulated by OGT. **A)** Overview of TGF-β signal transduction. Non-canonical signaling (ERK, p38, JNK) is shown on the left and canonical signaling (SMAD2/3) is shown on the right. **B)** Western blot (WB) for phosphorylated pan-Ser/Thr residues in HLFs. Cells were serum starved overnight followed by 2hr OSMI-1 pretreatment and or TGF-β1 stimulation (30min). **C)** WB for P-JNK and JNK in HLFs under same conditions as **A. D)** WB for P-ERK and ERK in HLFs under same conditions as **A. E)** WB for P-p38 and p38 in HLFs under same conditions as **A. F)** WB for P-p38 and p38 in HLFs pretreated with OSMI-1 for 2hrs followed by prolonged (15hr) stimulation with TGF-β1. Representative images are shown for each blot; densitometric quantification for each is indicated to the right. WBs are reported as fractional ratios of phospho/total protein and β-actin was utilized as a loading control. Data were analyzed with the two-way ANOVAs with Tukey-Kramer post-hoc comparisons tests; N=3-5 replicates per panel. All graphs are shown as mean ± SEM; *p<0.05, **p<0.01, ***p<0.001; ****p<0.0001.

### O-GlcNAc-modified p38 MAPK regulates p47^**phox**^ phosphorylation

Since blockade of O-GlcNAc transfer attenuated TGF-β-induced p38 MAPK phosphorylation, we posit that p38 MAPK might be O-GlcNAc-modified in HLFs. To enrich for this modification, we stimulated HLFs with TGF-β1 in the presence of O-GlcNAcase (OGA) inhibitor, Thiamet G^37^. Following immunoprecipitation (IP) of O-GlcNAc (via CD110.6), endogenous lysates were blotted for p38 MAPK, confirming the O-GlcNAcylation of p38 (**Fig 4a**). Interestingly, kinomic profiling of p38 MAPK-peptide substrates revealed robust changes in peptide phosphorylation in response to OGT inhibition (**Fig 4b**). These data provide additional evidence that TGF-β-induced p38 MAPK kinase activity is regulated by OGT activity. Among the p38 peptide substrate targets, OGT blockade most notably reduced the phosphorylation of the NCF1 (p47^phox^) peptide (**Fig 4c**), which likely reflects diminished p38 kinase activity as a consequence of lower O-GlcNAcylation.

**Figure 4.**
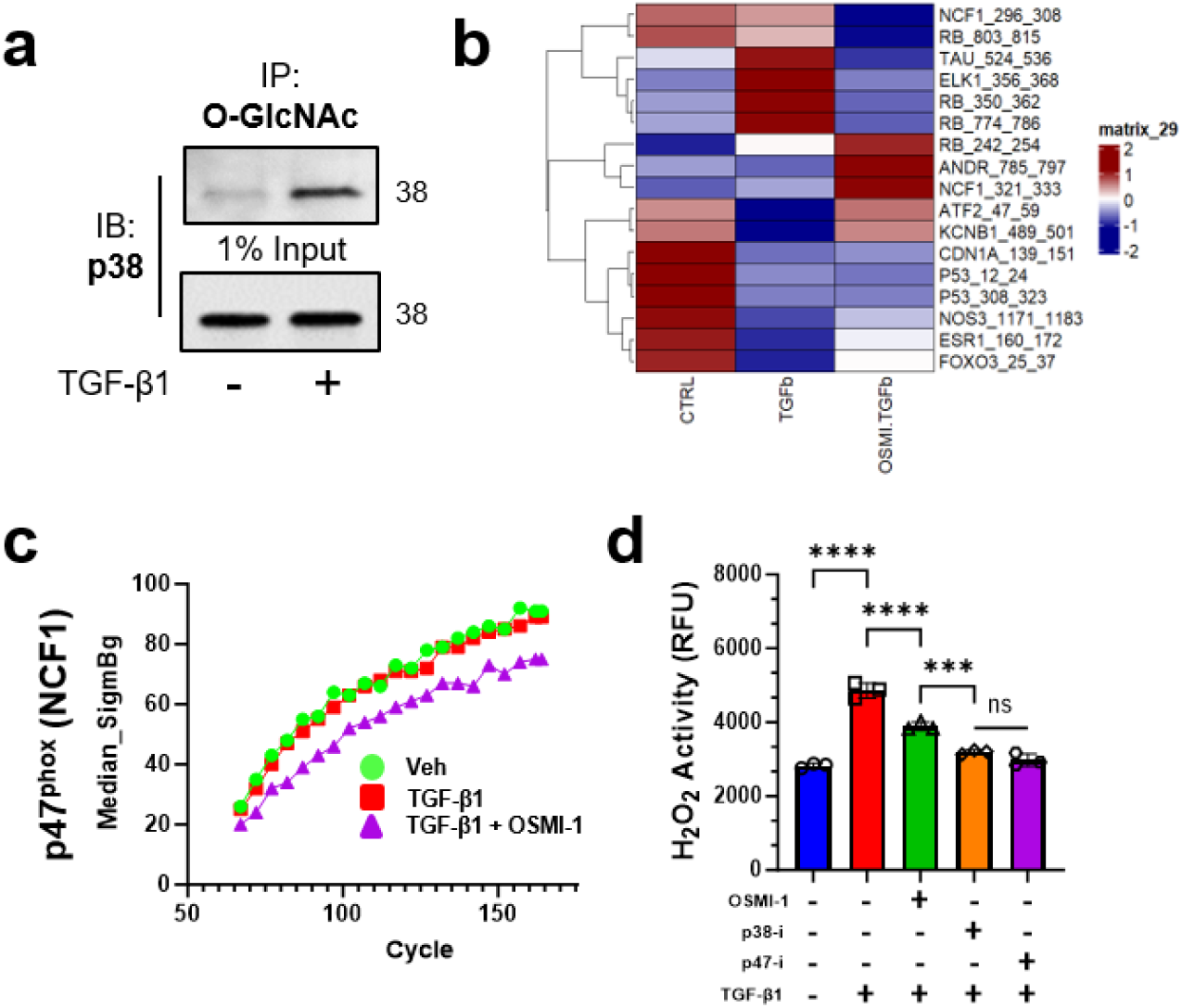
O-GlcNAc modified p38 MAPK promotes p47^**phox**^ phosphorylation and activity. **A)** Immunoblot for p38 following O-GlcNAc immunoprecipitation (IP) in HLFs. HLFs were treated overnight with Thiamet G (OGA inhibitor) and/or TGF-β1 prior to ionic-lysis and IP. A 1% input lysate is shown below. **B)** Heat map for predicted p38-modified peptides from HLF kinomics study. NCF1 (p47^phox^) phosphorylation was the most differentially regulated by TGF-β/OSMI-1 stimulation **C)** Dissimilar phosphorylation kinetic curves of p47^phox^ under the labeled conditions. Phosphorylation was measured by FITC intensity at each indicated cycle. **D)** H_2_O_2_ assay in HLFs. HFLs were serum starved overnight followed by 14-15 hour incubation with TGF-β1. H_2_O_2_ release was measured by the conversion of homovanillic acid to a fluorescent dimer (see *Methods*). p47inhibitor = 20μM; p38 inhibitor = 20μM. Data are reported as mean ± SEM and analyzed with one-way ANOVA; N=3 replicates per condition. ***p<0.001; ****p<0.0001

### TGF-β-induced ROS production is dependent on O-GlcNAc modified p38 MAPK

p47^phox^ is an essential subunit of the NOX2 reactive oxygen species (ROS)-generating complex. Since OGT is vital for phosphorylation of p47^phox^, we next sought to determine whether OGT inhibition results in a reduction in the TGF-β-induced ROS production in HLFs. The robust generation of ROS following TGF-β1 stimulation was markedly decreased by OGT inhibition (**Fig 4d**). Concomitantly, p38 and p47^phox^ pharmacologic inhibition independently resulted in an even greater reduction in ROS generation. Taken together with our IP data, these data support the notion that O-GlcNAc-modification of p38 is essential for the phosphorylation and activity of p47^phox^.

## iv. DISCUSSION

Idiopathic pulmonary fibrosis (IPF) is a devastating interstitial lung disease with poor prognosis and no cure. TGF-β-induced fibrogenesis is well described in the pathogenesis of IPF^38^. We previously discovered that the O-linked N-acetylglucosamine (O-GlcNAc) transferase (OGT) enhances canonical TGF-β-SMAD3 signaling in human IPF subjects and a murine pulmonary fibrosis disease model^33, 39^. Here, we demonstrate using an unbiased kinomics platform (**Fig 1**) that pharmacologic inhibition of OGT promotes widespread changes to kinase activity in human lung fibroblasts (HLFs) (**Fig 2a**). TGF-β1 treatment activated non-canonical TGF-β MAP kinases (p38, ERK, JNK) (**Fig 2b**), whereas loss of O-GlcNAc transfer prevented this response (**Fig 2c**). Molecular characterization revealed that TGF-β-induced phosphorylation of p38, but not ERK or JNK, was reduced by OGT inhibition (**Fig 3c-e**). TGF-β stimulation promoted p38 O-GlcNAcylation (**Fig 4a**), which was vital for the phosphorylation of the p38 substrate, p47^phox^, an integral regulator of cellular reactive oxygen species (ROS) (**Fig 4b-c**). Functionally, OGT inhibition attenuated TGF-β-induced ROS production in HLFs (**Fig 4d**), perhaps via loss of p38 O-GlcNAcylation. Taken together, this work couples unbiased kinomics with molecular validation to determine that pharmacologic inhibition of OGT induces widespread kinomic changes in HLFs. More specifically, we show that TGF-β1 stimulation promotes p38 MAPK O-GlcNAcylation, which enhances p47^phox^-induced ROS production in HLFs.

### O-GlcNAc and the pathogenesis of IPF

The O-GlcNAc modification is dependent on the availability of the OGT substrate, UDP-GlcNAc. Generation of this metabolic intermediate is dependent on the integration of carbohydrate, amino acid, fatty acid, and nucleotide metabolism. The O-GlcNAc modification therefore bridges cell signaling, function, and behavior with nutrient status. As metabolic dysfunction and reprogramming are well-described phenomena in the IPF lung, it is critical to understand whether the pathological manifestations of IPF are dependent on O-GlcNAcylation, yet to date little has been described. We previously reported that intratracheal instillation of OGT siRNA to the lung led to reversal of fibrosis in a non-resolving murine model of pulmonary fibrosis^33^. More specifically, we described that the O-GlcNAc modification is essential for the fibroblast-to-myofibroblast transition and collagen deposition/turnover in lung fibroblasts^33^. We also have shown that OGT inhibition controls the epigenetic regulation of anti-fibrotic genes, *Cox2* and *Homx1* even in the presence of TFG-β ^39^. While these studies provide a foundational understanding of O-GlcNAc biology in IPF, its functional and mechanistic roles in disease remain largely unexplored. Here, we demonstrate with unbiased kinomics that OGT blockade not only attenuates canonical TGF-β signaling but also promotes widespread kinomic changes (**Fig 2a**). Notably, several non-canonical kinases belonging to the MAPK family were activated by TGF-β but suppressed by OGT inhibition (**Fig 2,c**). In addition to the MAPK family, loss of O-GlcNAc transfer prevented TGF-β-activation of numerous cyclin dependent kinases, CDK3, CDK2, CDK4, CDK11, CDK5, CDK6, CDK9, CDK7 (**Fig 2b,c**). The O-GlcNAc modification has previously been shown to regulate CDK activity by blocking phosphorylation of upstream regulators (e.g. Cdc25, MYT1)^40^. Furthermore, O-GlcNAc is implicated in G1/S progression, replication, and mitosis; however, it is not yet known how these effects manifest in HLFs, particularly in the context of pulmonary fibrosis. As IPF fibroblasts exhibit dysregulated cell cycle control, including activation of CDK4/6-Rb-E2F signaling, increased G1/S progression, and altered expression of CDKs regulating DNA replication and mitosis, our findings suggest that the OGT/O-GlcNAc axis may contribute to dysregulated cell cycle control in IPF. Future studies will define the role of OGT in regulating CDK biology by examining how O-GlcNAcylation influences cell cycle progression in pulmonary fibrosis.

### O-GlcNAc-p38-p47^**phox**^ axis in human lung fibroblasts

Of the kinases activated by TGF-β stimulation, the MAPK family (ERK, JNK, p38) was enriched in our kinomics analysis. These kinases have been implicated in IPF and pre-clinical models of pulmonary fibrosis^19-21, 25, 27^. Here, we demonstrate that OGT blockade reduced TGF-β1-induced phosphorylation of p38 (**Fig 3e**). This effect was identified in an acute setting, maintained following prolonged stimulation (**Fig 3f**), and selective to p38 (not ERK or JNK) (**Fig 3c,d**). We also provide evidence that p38 phosphorylation at T180/Y182 is positively correlated with its O-GlcNAc modification (**Fig 3f, Fig 4a**). These data suggest that O-GlcNAc modification of p38 in HLFs is likely allosteric to T180 and/or Y182 and does not occupy the same site(s). It is also plausible that O-GlcNAc may modify and/or regulate the affinity of upstream kinase, MKK3/6 for p38. Both mechanisms independently may suggest that the O-GlcNAc modification is essential for p38 phosphorylation. To our knowledge, the O-GlcNAc modification of p38 has not been previously demonstrated in fibroblasts, and its role in fibrosis has not been studied, highlighting the novelty of the regulatory mechanism.

Downstream of p38, NCF1 (p47^phox^) was found to have differential phosphorylation between cells treated with TGF-β1 and those with TGF-β1 and OGT inhibition (**Fig 4c**). p47^phox^ functions in the assembly, membrane organization, and activation of the NADPH oxidase 2 (NOX2) complex, which is essential for the generation of reactive oxygen species (ROS)^41^. TGF-β signaling has been shown to upregulate NADPH oxidase components, including p47^phox^, supporting a role for p47^phox^-dependent ROS generation in fibroblast activation^42^. Although p47^phox^ expression has been reported in phagocytes and other cell types including HLFs^43^, its role in IPF remains incompletely understood. Loss of p47^phox^ has been shown to protect mice from bleomycin-induced lung injury, perhaps dependent on matrix metalloproteases^44^. Here, we further this mechanistic paradigm by implicating O-GlcNAc-modified p38 as central to p47^phox^ activation. Moreover, NOX2 expression has previously been observed in HLFs^45, 46^; however, NADPH oxidase 4 (NOX4) is vital for ROS generation contributing to the FMT and IPF pathogenesis^45, 47^. Given the expression patterns and functional homology between NOX2 and NOX4, it stands to reason that p47^phox^ may act on one or both homologs in HLFs. We provide evidence that inhibition of p47^phox^ completely abolishes TGF-β-induced H_2_O_2_ production in HLFs (**Fig 4d**) and loss of O-GlcNAc transfer impairs p47^phox^ phosphorylation (**Fig 4b,c**) and H_2_O_2_ production (**Fig 4d**). Taken together, these data suggest that p38-dependent regulation of p47^phox^ is essential for H_2_O_2_ production, which is controlled by OGT blockade. Ongoing work will determine the molecular mechanisms controlling p47^phox^-mediated H_2_O_2_ production, which is regulated by the O-GlcNAc modification in HLFs.

## Conclusions

Here, we show that pharmacologic inhibition of OGT disrupts non-canonical TGF-β signaling by suppressing activation of p38, ERK, and JNK in HLFs, as revealed by kinomic profiling. We demonstrate that p38 is O-GlcNAc modified in HLFs, and its phosphorylation and kinase activity are dependent on the O-GlcNAc modification. Activated p38-O-GlcNAc is a vital kinase of p47^phox^, which is essential for TGF-β-induced H_2_O_2_ production in HLFs. These findings identify O-GlcNAc transfer as a previously unrecognized regulator of non-canonical TGF-β signaling and redox biology in fibroblasts. Future studies are needed to evaluate the therapeutic benefit of modulating O-GlcNAc and/or OGT in the IPF lung.

## Limitations

This study utilizes an unbiased kinomics approach to predict kinase activity in HLFs. This approach is not without limitation as the PamArray® chips used in this study are restricted in the number of substrates available per chip, prohibiting a comprehensive analysis of kinase activity within cells. In addition, we exclusively assess acute TGF-β stimulated HLFs relative to controls. Although this approach limits cellular toxicity due to OGT inhibition with OSMI-1 and focuses on the exclusivity of TGF-β1 signaling, future studies are needed to confirm our findings in a prolonged/chronic system and evaluate the effects of additional growth factors on O-GlcNAc biology in HLFs.

## v. ACKNOWLEDGEMENTS

The authors declare financial support was received for the research, authorship, and/or publication of this article. NIH R01HL152246 to JWB; NIH R00HL131866 to JWB; NIH R01HL160911 to SK; NIH 5T32HD071866 to LIJ; NIH 5T32GM008361 to LIJ. We thank the UAB Center for Exercise Medicine and the UAB Medical Scientist (MD-PhD) Training Program for their financial and professional support.

## vi. METHODS

### Cell Culture

IMR-90 human lung fibroblasts (HLFs) were purchased from Coriell Institute and cultured as previously described^33^. Briefly, HLFs were cultured in Dulbecco’s Modified Eagle Medium supplemented with 10% Fetal Bovine Serum, 1X Penicillin-Streptomycin, and 1X GlutaMAX in a CO_2_ (5%) incubator at 37°C.

### Inhibitor treatment

OSMI-1 inhibitor was obtained from Sigma-Aldrich and reconstituted in DMSO to 25mM stocks that were used at a final concentration of 25µM for HLF treatment. HLF were pre-treated with OSMI-1 for 2 hours prior to 30 minutes of TGF-β1 (5ng/mL) stimulation.

### *In vitro* kinase assay

For kinomics assay, HLFs were lysed in MPER lysis buffer with 1X HALT inhibitors, 1X Z-PUGNAc, and Thiamet G. Kinomic profiling of lysates was performed by the UAB Kinome Core using standard methods as previously described. Briefly, protein lysates were quantified using a standard BCA method and 2 µg per sample was loaded for the serine/threonine (STK) array using the PamChip® (PamGene International’s-Hertogenbosch, The Netherlands). Phosphorylation data was collected over multiple computer-controlled pumping cycles and exposure times (10-200ms) for ∼144 substrates per array. Raw image analysis was conducted using Evolve2, and comparative analysis, upstream kinase predictions were done in BioNavigator v6.2 using the UpKin PamApp. After kinetic reads, post-wash captures were done at 10, 20, 50, 100, and 200ms exposures. These values were integrated into a slope, multiplied by 100 and Log2 transformed. Whole Chip comparative analysis (BioNavigator Upkin PTK v8.0/STK v6.0) was done between groups generating Mean Final Scores (MFS; non-directional combined specificity and sensitivity) and Mean Kinase Statistic (Direction of change with OSMI-1 or TGF-β1). Enrichment of each kinase in relation to the non-TGFβ stimulus is averaged based on all predicted kinase scoring from all peptides, creating a mean kinase statistic for each kinase.

### Unsupervised clustering analysis

Unsupervised hierarchical clustering of peptides (row) using Euclidian distance means-based hierarchal clustering method was used to generate a clustered heatmap using heatmap in R (V 4.5.0). Signal intensity (from peptide mean across all replicates) is colored from highest (red) to lowest (blue) intensity, and dendrogram trees annotate similarity clustering of peptides.

### Kinase prediction

Scored kinases from UpKin PamApp were plotted in GraphPad Prism to generate volcano plots comparing the effects of TGF-β1-induced kinase activity with and without OSMI-1 treatment. Alluvial plot demonstrating peptide substrates used for upstream kinase predictions was developed using the ggalluvial^48^ (V 0.12.6) extension of ggplot2 (V 4.0.2) in R (V 4.5.0).

### Western blotting

HLF cellular lysates were prepared for western blot as previously described ^33^. Briefly, isolated proteins were separated by SDS-PAGE using a 4-20% precast Ready Gel (Bio-Rad) and transferred to a 0.45 μm nitrocellulose membrane (Cytiva). Membranes were probed with the following primary antibodies: phosphoserine/threonine (PhosphoSolutions), p-ERK, ERK, p-p38, p38, p-JNK (Cell Signaling), JNK (R&D systems), and β-actin (Millapore Sigma). The following secondary antibodies were used: GtαMs IgG-HRP, GtαRb IgG-HRP, and RbαGt IgG-HRP (Thermo Fisher Scientific). All blots were imaged using the GE Imaging System (GE Healthcare, USA) and densitometric analyses was performed using FIJI software. All immunoblots were repeated for at least three independent trials with comparable results.

### Immunoprecipitation (IP)

Cells were lysed in RIPA buffer (Cell Signaling, #9806) supplemented with HALT protease and phosphatase inhibitors (ThermoScientific, 78442), DNAse 1 (IBI, XH28701) and Z-pugnac. A 1% lysate was saved for each sample as input material and prepared according to *Western Blotting*. Remaining lysates were precleared with protein L agarose beads, tumbling end-over-end for 1 hour at 4°C. Precleared lysates were incubated with anti-O-GlcNAc primary antibody (BioLegend #838004; CTD110.6; 4 μg) tumbling end-over-end at 4°C ON. The following day, protein L capture beads were added with end-over-end tumbling for 2 hours at 4°C. Beads were then washed 2X with ice-cold lysis buffer, 4X with detergent buffer (75mM Tris, 150mM NaCl, 1% Triton X-100), and 3X in 75mM Tris. Protein-bead complexes were dissociated at 95°C in 4X Laemmli + β-mercaptoethanol.

### H_2_O_2_ Assay

The release of extracellular hydrogen peroxide (H_2_O_2_) from adherent HLFs was quantified using a fluorometric homovanillic acid (HVA) oxidation assay, in which H_2_O_2_ reacts with HVA in the presence of horseradish peroxidase (HRP) to generate a fluorescent dimer. Cells were plated in 6-well dishes, serum-starved for 16 h, and stimulated with TGF-β1 (2.5 ng/mL) for 14–15 h prior to assay initiation. Cells were gently washed with HBSS and incubated for 2h at 37 °C in freshly prepared assay media containing 100 µM HVA and 5 U/mL HRP. Following incubation, assay media from each well was collected, mixed with NaOH/glycine/EDTA stop solution, and fluorescence was measured with a BioTek plate reader (excitation 321 nm, emission 421 nm). H_2_O_2_ concentrations were calculated from a standard curve.

### Quantification and statistical analysis

Statistical analysis description for each experiment can be found in the figure legends. Western blot images were quantified using FIJI and analyzed relative to β-actin. Statistical analyses were performed in Prism 10 (GraphPad). Error bars represent the mean ± standard error of the mean (SEM). A classic student t-test or one-way ANOVA with Tukey’s multiple comparison test was performed to assess significance. Statistical significance was represented as follows: *p<0.05, **p<0.01.

